# Microbial contamination in the genome of the domesticated olive

**DOI:** 10.1101/499541

**Authors:** Taylor Reiter, C. Titus Brown

**Affiliations:** Department of Population Health and Reproduction, University of California, Davis; Department of Food Science Graduate Group, University of California, Davis

## Abstract

Draft genomes of both the wild and domesticated olive were recently published. While working with these genomes, we identified contamination in the domesticated olive genome representing .06% of basepairs and 1.3% of scaffolds. We used targeted and untargeted approaches to identify the contaminating sequences, the majority of which were *Aureobasidium pullulans*. We applied the same method to the wild olive genome and did not find evidence of contamination. Although contamination in the domesticated olive genome was not prolific, it could lead to biased results either from functional content encoded in the contaminant sequences, or from using it to separate microbial reads from olive reads in microbiome studies.

## Introduction

Olives are agriculturally, nutritionally, economically, and socially important fruits produced from the tree *Olea europaea* var. *europaea*. The olive was domesticated approximately 6000 years ago in the Levant and is now cultivated outside of its original wild and domestic footprint (Besnard 2016). As with other domesticated species, the domesticated olive bears strong resemblance to its wild progenitor oleaster (*Olea europaea* var. *sylvestris*) (Unver et al. 2017).

Recently, draft genomes were published for both the domesticated olive and the wild olive. Both genomes contain 23 diploid chromosomes and an estimated size between 1.38-1.48 GB of sequence (Cruz et al. 2016; Unver et al. 2017). The domesticated olive genome (sequenced approximately one year prior to the wild olive genome) has been used for gene identification (Haberman et al. 2017), discovery of olive fruit development programs (D’Angeli and Altamura 2016), and to improve understanding of genome structure (Barghini et al. 2017).

While working with the domesticated and wild olive genomes, we found sequence consistent with contamination in the domesticated olive genome. Approximately .06% of basepairs were identified as non-plant sequence. We did not find contamination in the wild olive genome. The contamination in the domesticated olive genome is not a prolific problem, and contamination on this level could be common for many eukaryotic genomes (Delmont and Eren 2016). However, for experiments involving the olive microbiome, leaving this sequence in the genome could lead to biased results. We characterized the contaminant sequences, and suggest that these sequences be removed from the genome in subsequent analyses.

## Results and Discussion

### Identification of *Aureobasidium pullulans* contamination

The *A. pullulans* var. *santander* genome was published alongside the domesticated olive genome when the authors realized they had deeply sequenced an *A. pullulans* genome (Cruz et al. 2016). They mapped their reads to four *A. pullulans* genomes and used mapped reads to produce a partial draft genome of *A. pullulans* var. *santander*. However, we found *A. pullulans* sequence was still present in the final genome assembly.

To locate the scaffolds containing putative *A. pullulans* sequence, we aligned the domesticated olive genome against two *A. pullulans* genomes.

We first aligned to the *A. pullulans* var. *santander* genome, producing 140 alignments greater than 500 base pairs in length and with greater than 98 percent identity. The alignments were distributed across 65 scaffolds with an average length of 2,273 basepairs (minimum 534 bp, maximum 9,966 bp).

Given that the *A. pullulans* var. *santander* genome was only 2/3 complete (Cruz et al. 2016), we also aligned the domesticated olive genome to the *A. pullulans* variant *EXF-150* genome as it contained additional sequence. We found 176 alignments greater than 500 base pairs in length with greater than 94 percent identity. The average length was 1,648 basepairs (minimum 556, maximum 10,548).

Alignments with the *A. pullulans* var. *EXF-150* genome were shorter on average than those for the *A. pullulans* var. *santander* genome. Because the var. *santander* genome represents the exact genome of the contaminant present in the domesticated olive genome, these alignments were likely longer due to less strain variation. However, there were more matches with the complete *A. pullulans* var. *EXF-150* genome than with the *A. pullulans* var. *santander* genome.

When ranges from both alignments were combined, 49 domesticated olive genome scaffolds aligned completely to *A. pullulans* sequence. An additional 59 scaffolds had partial, high-sequence identity alignments.

### Identification of other potential contamination

We also pursued a non-specific approach to identify other contaminants. We calculated tetranucleotide frequency for each of the 11,038 scaffolds in the domesticated olive genome, and performed pairwise comparisons for each scaffold (*Figure 1*). We calculated the average similarity for each scaffold and selected a similarity cutoff of two standard deviations below the mean similarity as an approximate identifier for outliers. Using this criteria, we identified 463 outlier scaffolds.

**Figure 1:**
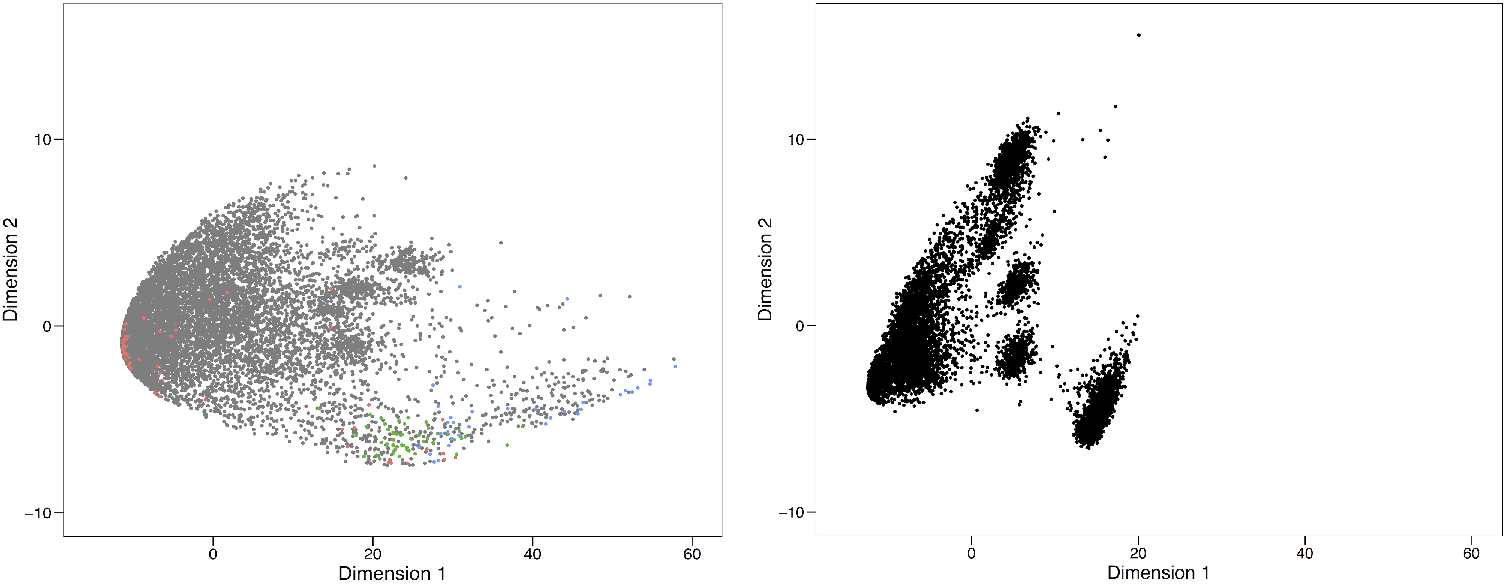
MDS plots of the domesticated olive genome (left) and the wild olive genome (right). Each point represents a contiguous sequence in the genome, and the distance between each point represents similarity. The domesticated olive genome is more dispersed than the wild olive genome. Additionally, colored points indicate contiguous sequences identified as contamination in the domesticated olive genome. Red points correspond to chimeric *Aureobasidium pullulans* scaffolds, green points correspond to non-chimeric *A. pullulans* scaffolds, and blue corresponds to contaminants identified by BLAST. Grey points represent non-contaminant contiguous sequences. Many chimeric scaffolds do not separate from the core genome contiguous sequences, suggesting they do contain real olive sequence. Other contaminants clearly separate.

We further classified these outliers by BLASTing them against the NCBI’s nt database. Using BLAST matches, we filtered these outliers to avoid false positives. We removed any sequence with no BLAST match (263 scaffolds), any scaffold that matched repetitive sequences in the genus *Olea* (89 scaffolds), any scaffold where the best BLAST match had a bitscore lower than 100 (59 scaffolds), any scaffold where the best BLAST match or the majority of BLAST matches were to another plant (7 scaffolds), and scaffolds where the only BLAST match for a contig had a low percent identity (< 80%) and a small length (< 259) (3 scaffolds).

We identified 42 contaminant scaffolds using this method. However, nine scaffolds overlapped with those previously identified as *A. pullulans*. Therefore, we identified a total of 33 contaminant scaffolds with BLAST, totalling 461,345 base pairs and with an average length of 13,980 (Table 1).

**Table 1.**
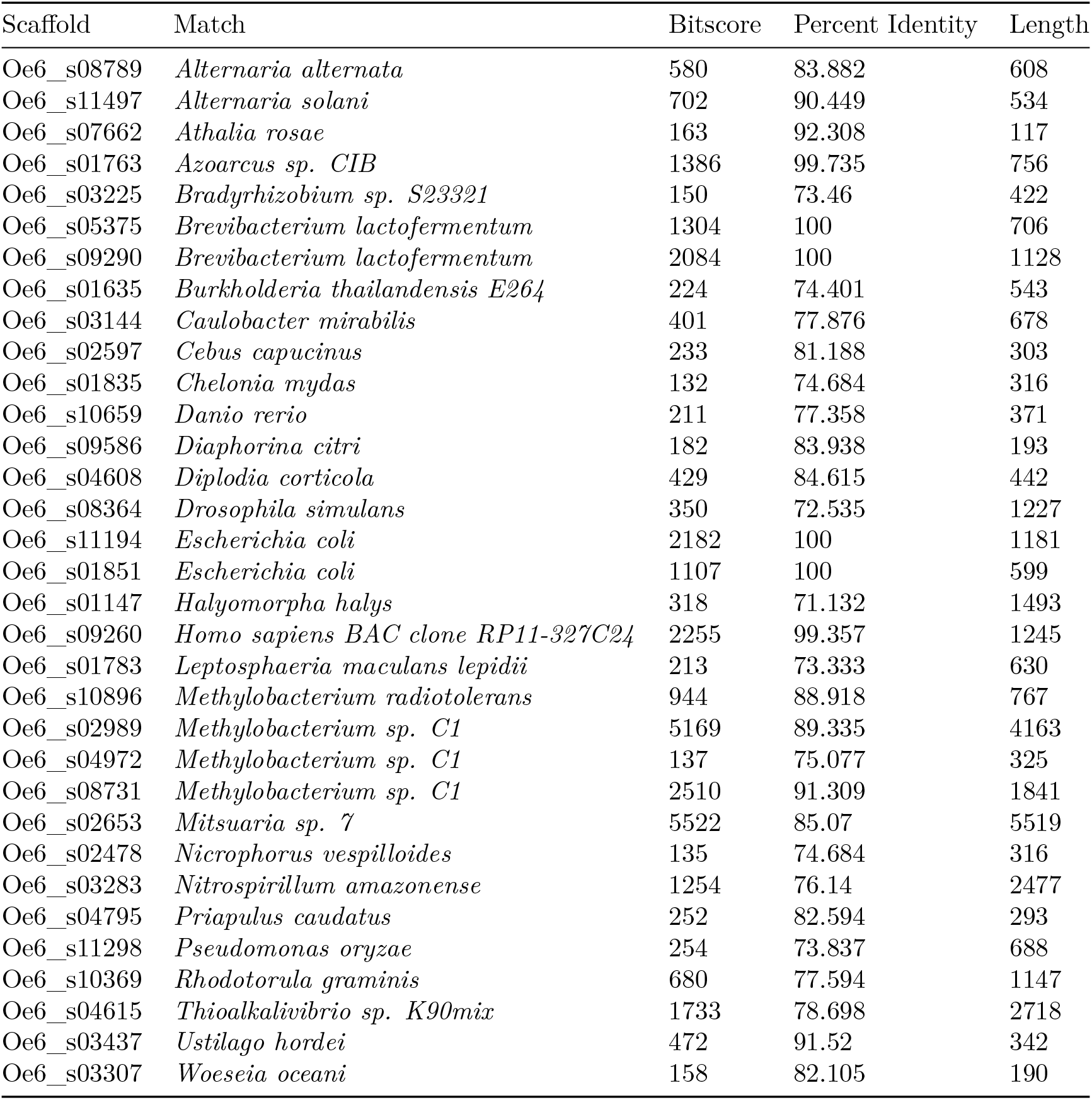
Domesticated olive BLAST matches.

### Screening of the wild olive genome for contamination

We performed the same tetranucleotide frequency clustering and BLAST analysis on the wild olive genome. Two hundred and fifty eight potential contaminants were identified. However, almost all had strong BLAST matches to other plant species. We noted only four scaffolds with a significant BLAST match to sequence not of plant origin (Table 2). None of these scaffolds were assembled into the 23 chromosomes in the final wild olive genome assembly, but remain in the genome as unplaced scaffolds.

**Table 2.**
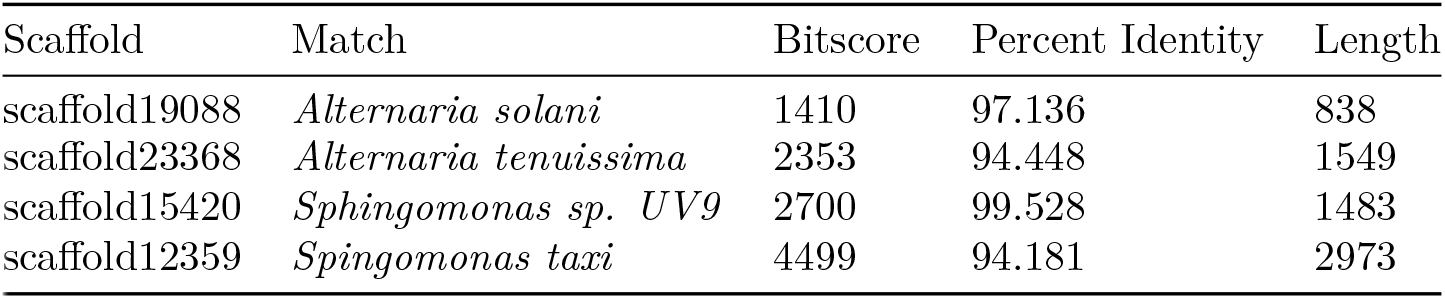
Wild olive genome BLAST matches

### Evaluating the likelihood of horizontal gene transfer versus contamination

We reasoned that if the non-plant origin nucleotide sequences that we saw in either the domesticated and wild olive genomes were present in only one genome, then the nucleotide sequences were likely contamination. If they were present in both genomes, then the sequences could represent horizontal gene transfer that occurred in the *Olea europaea* lineage before the split between the last common ancestor of the domesticated and wild olive species.

To evaluate whether sequence ranges identified as potential contaminants in one olive genome were present in the other olive genome, we performed whole genome alignment of the domesticated and wild olive genomes. We first evaluated whether any sequence characterized as *A. pullulans* in the domesticated olive genome was present in the wild olive genome. No scaffold that exclusively contained *A. pullulans* sequence aligned to the wild olive genome. Of the scaffolds that were characterized as chimeric (e.g. contained both *A. pullulans* and olive sequence), six ranges of *A. pullulans* sequence overlapped with the wild olive genome. When we investigated these sequences further, they were sparsely populated with real nucleotides and primarily contained ambiguous characters (e.g. “N”). We concluded that *A. pullulans* sequence was uniquely contained in the domesticated olive genome.

No identical contaminant BLAST matches were in both the domesticated and wild olive genome. However, we also checked whether the nucleotide ranges identified as microorganisms in our BLAST matches aligned to the other olive genome.

None of the 33 domesticated olive genome aligned contaminant scaffolds aligned to the wild olive genome. Of the four wild olive genome contaminant scaffolds, only one aligned to the domesticated olive genome. Scaffold scaffold19088, which had the highest BLAST match to *Alternaria solani*, aligned to the middle of a large scaffold in the domesticated olive genome (nucleotides 1150907-1150149, scaffold length 1326173). Therefore, we posit that this BLAST match may be evidence of horizontal gene transfer in both olive genomes. We did not perform further tests to evaluate the liklihood of a horizontal gene transfer event (e.g. site rate heterogeneity).

Interestingly, this nucleotide sequence was not detected by tetranucleotide frequency clustering in the domesticated olive genome, indicating that the method we used for identification of nonspecific contamination is only robust for scaffolds in which the contaminant sequence skews the tetranucleotide abundance of the entire scaffold. This leaves the possibility that small contaminant sequences accidentally assembled into larger olive sequences (e.g. chimeric scaffolds) may still be present in both olive genomes.

We recognize that whole genome alignment would not detect horizontal gene transfer events that occurred after the split between the last common ancestor of the domesticated and wild olive species (e.g., in the last 6000 years). However, given the rarity of horizontal gene transfer events between plants and microorganisms (Richards et al. 2009), we assume that the majority of non-plant origin sequence found in the domesticated and wild olive genomes is contamination.

### Additional functional potential encoded in the domesticated olive genome

We were curious whether the contaminant sequences in the domesticated olive genome conferred increased functional potential upon the genome. We compared ranges of sequences we identified as *A. pullulans* against the domesticated olive genome annotations. We found 94 predicted proteins that were attributable to *A. pullulans* sequence. We BLASTed a subset of these proteins against the NCBI nr database and found that all matched *A. pullulans* proteins, and the majority matched *A. pullulans* hypothetical or predicted proteins.

We then performed a similar search for ranges that overlapped between the BLAST matches and those in the annotation file. One hundred eighty three protein coding genes were predicted on 33 scaffolds. We BLASTed a subset of these protein sequences against the NCBI nr database to confirm the contaminant identity and to identify the function of the predicted protein. For example, a transcript encoded on scaffold 0e6_s01635 coded for product 0E6A107529P1. The protein product matched terminase large subunit from the phylum *Proteobacteria*. Another transcript encoded on scaffold 0e6_s02653 coded for product 0E6A064969P1, which matched serine/threonine protein phosphatase, also from the phylum *Proteobacteria*. These sequences could be of mitochondrial or chloroplast origin, however sequences matching mitochondria or chloroplast were filtered from the domesticated olive genome (Cruz et al. 2016)

Taken together, these results suggest that the contamination in the domesticated olive genome has misrepresented the functional potential of the genome.

## Conclusion

We identified contamination in the domesticated olive genome representing .06% of basepairs and impacting 1.3% of scaffolds. The majority of contamination is attributable to *A. pullulans*. As some of these sequences code for proteins, the functional potential of the domesticated olive genome is currently misrepresented. We found little evidence for contamination in the wild olive genome. We recommend that contaminant sequences be removed from the domesticated olive genome assembly for subsequent analyses.

## Methods

All code used in these analyses is available at https://github.com/taylorreiter/olive_genome. All files output by our analyses, including the contaminant-masked domesticated olive genome, are available at https://osf.io/yj2cu/. All associated files ares available for download there as well, save for the comparison matrices as there were multi-gigabyte files and were too large to store on OSF. However, instructions for repeating the analysis and re-generating all files are available in the github repository.

### Data

We downloaded the domesticated olive genome and accompanying files from http://denovo.cnag.cat/genomes/olive/. All analyses were completed with the Oe6 genome. We downloaded the wild olive genome and accompanying files from http://olivegenome.org/downloads/. We used Olea_europaea>1Kb_scaffolds.fa due to the large number of scaffolds in the full genome (over 2 million).

### Whole genome alignments

We used the `nucmer` function in the Mummer software package to produce alignments between *A. pullulans* genomes and the domesticated olive genome, and the domesticated olive genome and wild olive genome (Kurtz et al. 2004). We used default parameters. For *A. pullulans* alignments, we filtered results below 500 base pairs. For the domesticated and wild olive alignment, we filtered results below 100 base pairs.

### Contaminant identification

We used the `compute` function in the sourmash software package to calculate tetramernucleotide frequency signatures for each scaffold in the domesticated and wild olive genomes (Brown and Irber 2016). We used a k-mer size of 4, a scaled value of 1, and tracked abundance of each k-mer. We performed pairwise comparison of these signatures with the `compare` function in the sourmash software package. We BLASTed scaffolds with a similarity to all other scaffolds two standard deviations below the mean scaffold similarity against the NCBI nt database using `blastn` (Altschul et al. 1990). The NCBI database was downloaded on 2/27/2018.

### Contaminant removal

We masked the contaminants we identified in the Oe6 genome using bedtools (Quinlan 2014). For contaminant scaffolds identified with BLAST and contaminant scaffolds that completely aligned to *A. pullulans*, we removed full scaffolds. For contaminant scaffolds that partially aligned to *A. pullulans*, we masked partial scaffolds.

## References

Altschul Stephen F, Warren Gish, Webb Miller, Eugene W Myers, and David J Lipman. 1990. “Basic Local Alignment Search Tool.” Journal of Molecular Biology 215 (3). Elsevier: 403–10.

Barghini Elena, Flavia Mascagni, Lucia Natali, Tommaso Giordani, and Andrea Cavallini. 2017. “Identification and Characterisation of Short Interspersed Nuclear Elements in the Olive Tree (Olea Europaea L.) Genome.” Molecular Genetics and Genomics 292 (1). Springer: 53–61.

Besnard Guillaume. 2016. “Origin and Domestication.” In The Olive Tree Genome, 1–12. Springer.

Brown C. Titus, and Luiz Irber. 2016. “Sourmash: A Library for MinHash Sketching of DNA.” The Journal of Open Source Software 1 (5). The Open Journal: 27.https://doi.org/10.21105/joss.00027.

Cruz Fernando, Irene Julca, Jèssica Gómez-Garrido, Damian Loska, Marina Marcet-Houben, Emilio Cano, Beatriz Galán, et al. 2016. “Genome Sequence of the Olive Tree, Olea Europaea.” GigaScience 5. Oxford University Press: 29–29.

D’Angeli Simone, and Maria Maddalena Altamura. 2016. “Unsaturated Lipids Change in Olive Tree Drupe and Seed During Fruit Development and in Response to Cold-Stress and Acclimation.” International Journal of Molecular Sciences 17 (11). Multidisciplinary Digital Publishing Institute: 1889.

Delmont Tom O, and A Murat Eren. 2016. “Identifying Contamination with Advanced Visualization and Analysis Practices: Metagenomic Approaches for Eukaryotic Genome Assemblies.” PeerJ 4. PeerJ Inc.: e1839.

Haberman Amnon, Ortal Bakhshian, Sergio Cerezo-Medina, Judith Paltiel, Chen Adler, Giora Ben-Ari, Jose Angel Mercado, Fernando Pliego-Alfaro, Shimon Lavee, and Alon Samach. 2017. “A Possible Role for Flowering Locus T-Encoding Genes in Interpreting Environmental and Internal Cues Affecting Olive (Olea Europaea L.) Flower Induction.” Plant, Cell & Environment. Wiley Online Library.

Kurtz Stefan, Adam Phillippy, Arthur L Delcher, Michael Smoot, Martin Shumway, Corina Antonescu, and Steven L Salzberg. 2004. Genome Biology 5 (2). Springer Nature: R12. https://doi.org/10.1186/gb-2004-5-2-r12.

Quinlan Aaron R. 2014. “BEDTools: The Swiss-Army Tool for Genome Feature Analysis.” Current Protocols in Bioinformatics. Wiley Online Library, 11–12.

Richards Thomas A, Darren M Soanes, Peter G Foster, Guy Leonard, Christopher R Thornton, and Nicholas J Talbot. 2009. “Phylogenomic Analysis Demonstrates a Pattern of Rare and Ancient Horizontal Gene Transfer Between Plants and Fungi.” The Plant Cell 21 (7). Am Soc Plant Biol: 1897–1911.

Unver Turgay, Zhangyan Wu, Lieven Sterck, Mine Turktas, Rolf Lohaus, Zhen Li, Ming Yang, et al. 2017. “Genome of Wild Olive and the Evolution of Oil Biosynthesis.” Proceedings of the National Academy of Sciences 114 (44). National Acad Sciences: E9413–E9422.

